# Lipocalin 2 dictates cancer cell plasticity elicited by therapy-induced senescence

**DOI:** 10.1101/2022.03.08.483463

**Authors:** Jorge Morales-Valencia, Lena Lau, Teresa Martí-Nin, Ugur Ozerdem, Gregory David

**Affiliations:** Department of Biochemistry and Molecular Pharmacology, NYU Langone Medical Center, New York, NY 10016, USA; NYU Langone Perlmutter Cancer Center, NYU Langone Health, New York, NY 10016, USA; Department of Pathology, New York University School of Medicine, NYU Langone Health, New York, NY 10016, USA; Department of Urology, New York University School of Medicine, NYU Langone Health, New York, NY 10016, USA

## Abstract

The acquisition of novel detrimental cellular properties following exposure to cytotoxic drugs leads to aggressive and metastatic tumors that often translates into an incurable disease. While the bulk of the primary tumor is eliminated upon exposure to chemotherapeutic treatment, residual cancer cells and non-transformed cells within the host can engage a stable cell cycle exit program named senescence. Senescent cells secrete a distinct set of pro-inflammatory factors, collectively termed the senescence-associated secretory phenotype (SASP). Upon exposure to the SASP, cancer cells undergo cellular plasticity resulting in increased proliferation, migration and epithelial-to-mesenchymal transition. The molecular mechanisms by which the SASP regulates these pro-tumorigenic features are poorly understood. Here, we report that breast cancer cells exposed to the SASP strongly upregulate Lipocalin 2 (LCN2). Furthermore, we demonstrate that LCN2 is critical for SASP-induced increased migration in breast cancer cells, and its inactivation potentiates the response to chemotherapeutic treatment in mouse models of breast cancer. Finally, we show that neoadjuvant chemotherapy treatment leads to LCN2 upregulation in residual human breast tumors, and correlates with worse overall survival. These findings provide the foundation for targeting LCN2 as an adjuvant therapeutic approach to prevent the emergence of aggressive tumors following chemotherapy.

## Introduction

Breast cancer affects more than one in ten women worldwide (1). Currently, neoadjuvant chemotherapy is extensively used to treat breast cancer patients as it reduces tumor burden, thus downstaging the disease (2). However, most non-targeted anticancer agents do not only trigger cytotoxicity in dividing cells, but also engage specific cellular response programs, including senescence, on both cancer cells and the tumor microenvironment (3).

Cellular senescence refers to the stable cell proliferation arrest caused by either telomere shortening, oncogene activation or genotoxic stress, all of which converge towards the activation of a sustained DNA damage response (DDR) (4). Because of its engagement in preneoplastic lesions, senescence was originally hypothesized to serve as a barrier to malignant transformation, by preventing the proliferation of cells harboring an altered genetic content (5). Additionally, senescent cells accumulate over time in mammals and contribute to the health defects associated with aging (6,7). The beneficial impact of senescence as a tumor suppressor mechanism early in life along with its detrimental impact on aging phenotypes led to the theory of antagonistic pleiotropy of senescence (8). One of the hallmarks of senescence that could rationalize these otherwise contradictory features is the senescence-associated secretory phenotype (SASP) (9). The SASP consists of a discrete set of pro-inflammatory cytokines, chemokines and growth factors secreted by senescent cells, in a cellular and senescence inducer-specific manner. As such, the SASP may contribute to the “inflamm-aging”, a sterile inflammation that develops as individuals age. In addition to its potential impact on aging phenotypes, exposure to the SASP has been reported to promote aggressive traits in tumor models, including increased cellular proliferation (10), enhanced angiogenesis (11) and activation of the epithelial-to-mesenchymal transition (EMT) (12). However, the molecular mechanisms engaged by senescent cells to drive tumorigenesis remain poorly understood.

Lipocalin 2 (LCN2), also referred to as neutrophil gelatinase-associated lipocalin (NGAL), is a 25 k-Da secreted glycoprotein involved in iron metabolism and inflammation (13). LCN2 protein expression levels are increased in various cancer types including breast (14, 15), colon (16) and pancreatic (17). Accordingly, high LCN2 expression has been detected in carcinoma tissues, sera and urine of breast cancer patients (18).

Here, we demonstrate that exposure to the SASP or senescence-inducing neoadjuvant chemotherapy results in the potent upregulation of LCN2 expression in breast cancer cells *in vitro* and in human cancer samples, respectively, which correlates with increased cellular plasticity and poor prognosis.

## Results

### The IL-1-dependent SASP promotes cellular plasticity in breast cancer cells

We have previously demonstrated that inactivation of the IL-1 pathway can be used to uncouple SASP production from senescence-associated cell cycle exit (19). We leveraged this property of IL-1α-inactivated cells to determine the impact of SASP exposure on cancer cells’ properties. MCF7 breast cancer cells were exposed to conditioned media (CM) from wild-type (WT) TERT-immortalized IMR90 (IMR90T) fibroblasts, WT IMR90Ts rendered senescent through ectopic expression of oncogenic Ras^G12V^, and senescent IL-1α^-/-^ IMR90Ts (Figure 1A). MCF7 cells exposed to CM from senescent WT cells (RasCM) migrated significantly faster than cells exposed to CM from growing cells in a scratch assay (Figure 1B, C). Strikingly, CM collected from senescence IL-1α^-/-^ cells was unable to promote cancer cell migration (Figure 1B, C). These observations suggest that exposure to the IL-1α-dependent SASP is sufficient to stimulate breast cancer cells’ migration. To determine whether SASP-induced increased migration correlates with enhanced chemotactic capacities, MCF7 cells cultured with CM for 2 days were allowed to migrate for 48 hours in a transwell assay. MCF7 cells cultured with Ras CM displayed increased transwell migration compared to cells cultured with growing CM or cells cultured with CM from IL-1α^-/-^ senescent cells (Figure 1D). Consistent with our previous demonstration that SASP production is dependent on the IL-1α/IL-1R axis, chemotactic-based migration was significantly reduced when CM was obtained from senescent IMR90Ts depleted for IL-1R (Figure 1E). Furthermore, exposure to senescent CM resulted in morphology changes in MCF-7 cells, which then adopted a fibroblast-like appearance, a feature of EMT (20). By contrast, MCF7 cells treated with CM from senescent IL-1α^-/-^ cells maintained their cobblestone-like morphology and strong cell-cell adhesions (Figure 1F). Accordingly, MCF7 cells exposed to senescent CM exhibited loss of expression of the epithelial marker E-cadherin (Figure 1G). Finally, the proportion of MCF7 cells expressing the EMT-associated surface marker CD44 (21) increased upon exposure to SASP (Figure 1H). Taken together, these results indicate that exposure to the IL-1-dependent SASP induces cellular plasticity in breast cancer cells, as evidenced by increased migratory properties and the engagement of an at least a partial EMT program.

**Figure 1.**
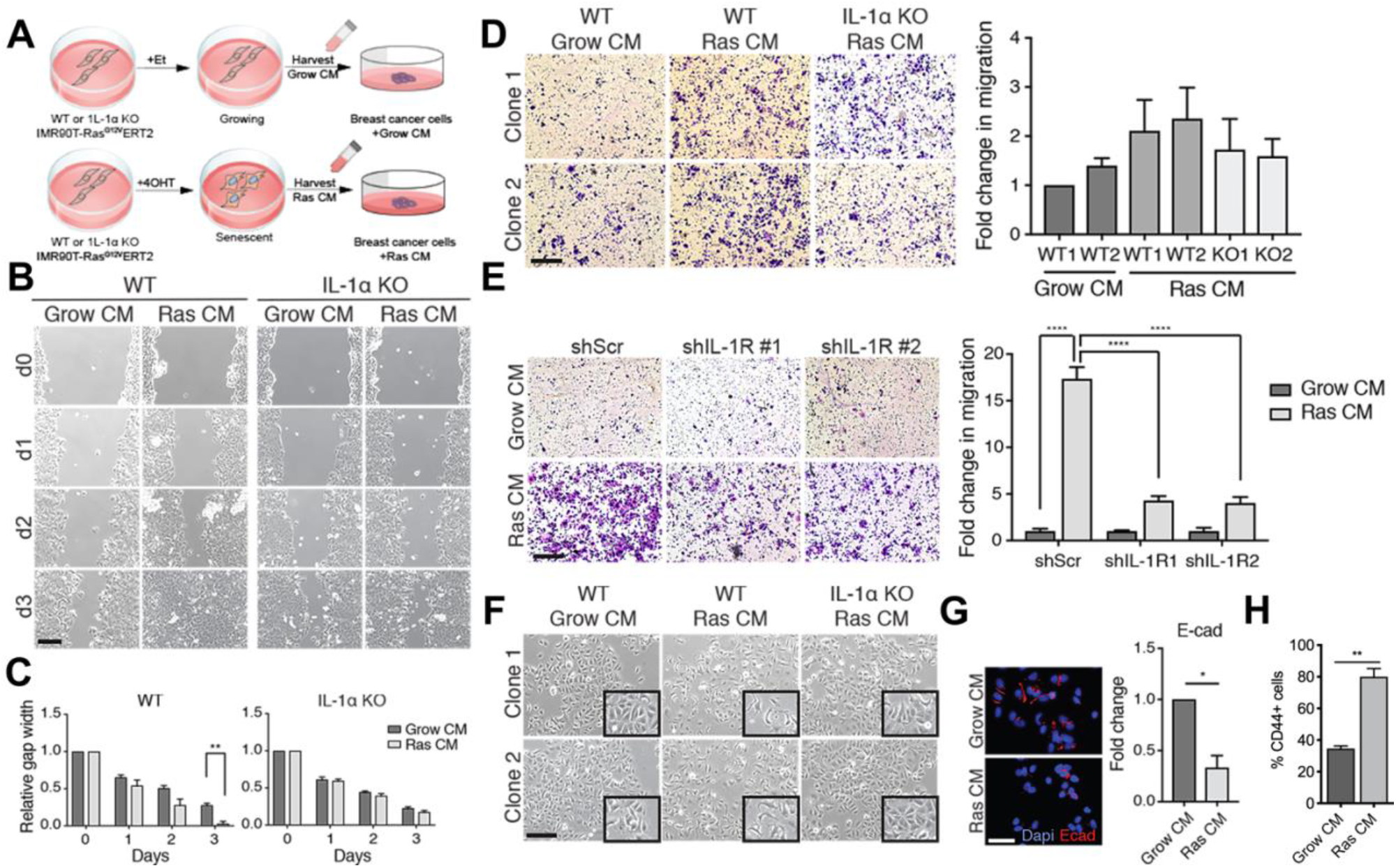
The IL-1-dependent SASP promotes cellular plasticity in breast cancer cells. (A) MCF7 breast cancer cells were exposed to conditioned media (CM) from wild-type (WT) TERT-immortalized IMR90 (IMR90T) fibroblasts, senescent WT IMR90Ts (through ectopic expression of oncogenic Ras, Ras^V12^) and senescent IL-1α^-/-^ IMR90Ts (B) Representative images of scratched MCF7 cells cultured in conditioned media (CM) from growing (Grow CM) or Ras-senescent (Ras CM) IMR90T cells with or without IL-1α for 3 days. Scale bar = 200 um (C) Quantification of relative gap width of MCF7 cultures treated with the indicated CM. (D, E) Representative images and quantification of transwell migration assays of MCF7 cells treated with the indicated CM. (F) Representative images of MCF7 cells treated with CM from 1 of 2 IMR90T clones, either growing or Ras-induced senescent with or without IL-1α. Inset: magnified images. Scale bar = 200 um. (G) Representative images and quantification of E-cadherin immunofluorescence staining in MCF7 cells treated with the indicated CM. Scale bar = 50 um. (H) Quantification of CD44^+^ MCF7 cells after being treated with the indicated CM as analyzed by flow cytometry. n = 3, * p < 0.05, ** p < 0.01.

### Exposure to the SASP induces expression of Lipocalin-2 in breast cancer cells

To begin to decipher the molecular mechanisms underlying the impact of exposure to the SASP on breast cancer cells’ properties, we profiled the transcriptome of MCF7 cells exposed to CM from growing or senescent IMR90Ts. Using a log2 fold change cutoff 1 and an adjusted p-value of < 0.05, 1981 genes were found differentially expressed between MCF7 cells exposed to growing CM and MCF7 exposed to senescent CM. Pathways upregulated in senescent CM samples include inflammatory response and extracellular matrix organization (Figure 2A). Gene sets that were significantly enriched in senescent CM compared to growing CM samples included Epithelial to Mesenchymal Transition and Protein Secretion (Figure 2B). These results are consistent with the previous demonstration that exposure to the SASP activates an inflammatory response and the initiation of an EMT program (22, 23). The most upregulated transcript in MCF7 exposed to senescent CM encodes the protein Lipocalin-2 (LCN2, or NGAL) (Figure 2C). We confirmed the upregulation of LCN2 mRNA and protein levels via qRT-PCR (Figure 2D) and Western Blot (Figure 2E). LCN2 was not upregulated in MCF7 cells treated with CM from senescent IL-1α^-/-^ cells (Figure 2D, E). Exposure to SASP from cells rendered senescent by etoposide (Etop) treatment resulted in a significant increase in LCN2 mRNA levels, indicating that LCN2 upregulation was independent of the stimulus used to induce senescence (Figure 2F). Of note, the LCN2 upregulation induced by exposure to genotoxic stress-induced SASP also correlated with increased migration (Figure 2G). We extrapolated these observations to independent breast cancer cells with various ER, PR or HER2 status, and detected a consistent upregulation of LCN2 upon exposure to the SASP (Figure 2H). In these conditions, these cells also exhibited an elongated and mesenchymal-like morphology (data not shown). In addition, we observed a significant increase in the rate of migration of MDA-MB-231 cells exposed to the SASP for 2 days (Figure 2I). Taken all together, these results indicate that the SASP secreted from cells induced to senesce by various stimuli promotes migration and LCN2 upregulation in several breast cancer cell lines.

**Figure 2.**
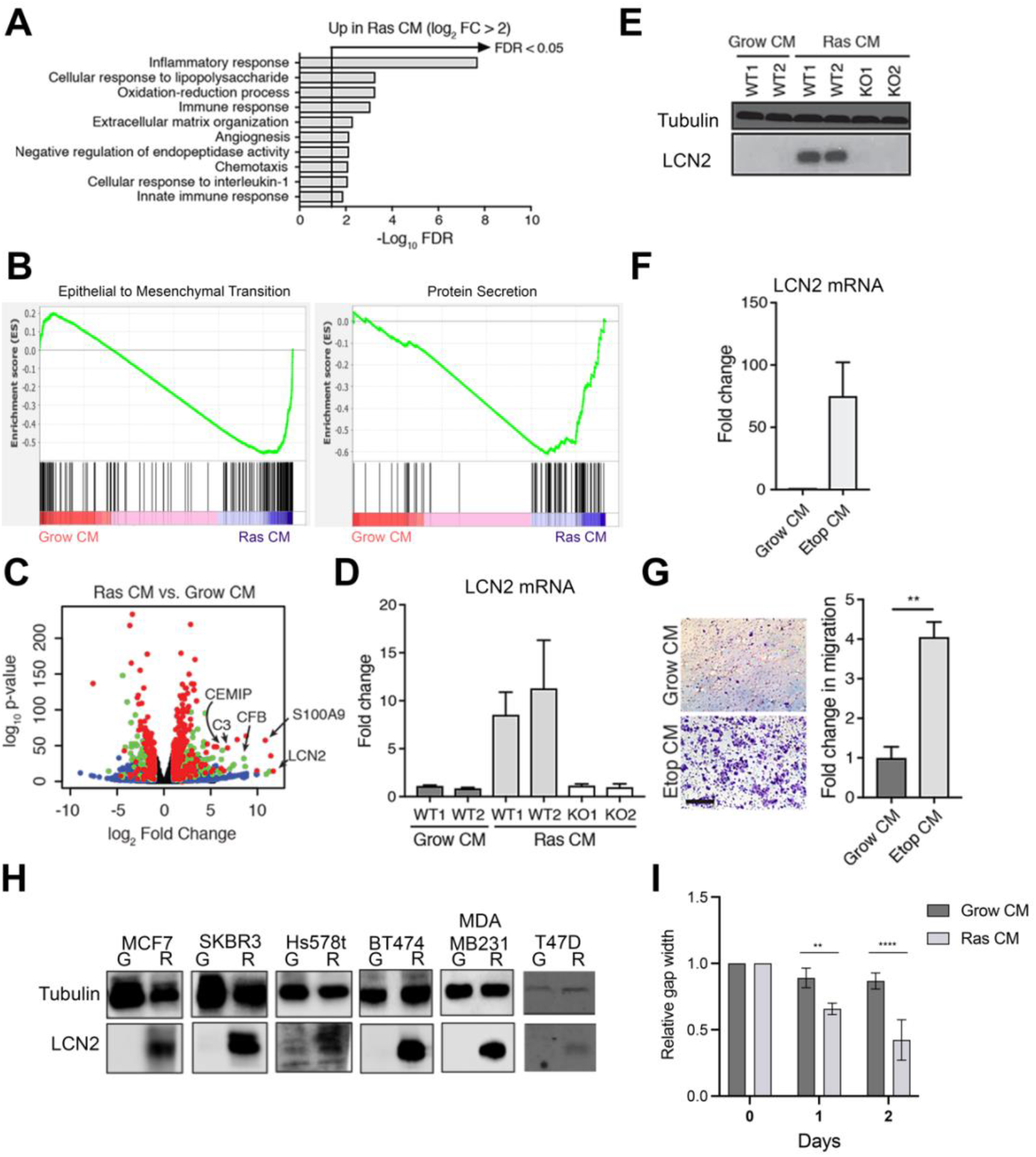
Exposure to the SASP induces expression of LCN2 in breast cancer cells. (A) Gene ontology analysis of genes that are upregulated in Ras CM compared to Grow CM samples with a log2 fold change > 2. (B) GSEA plots of pathways upregulated in Ras CM compared to Grow CM samples. (C) Volcano plot depicting differentially expressed genes in Ras CM samples compared to Grow CM samples. (D) qRT-PCR analysis and (E) Western blot for LCN2 expression in MCF7 cells cultured with the indicated CM. (F) qRT-PCR analysis for LCN2 expression in MCF7 cells cultured with Grow CM or conditioned media from etoposide-induced senescent IMR90T cells (Etop CM) (G) Representative images and quantification of transwell migration assay of MCF7 cells treated with the indicated CM. (H) Western blot for LCN2 expression in MCF7, SKBR3, Hs578t, BT474, MDA-MB-231 and T47D cells cultured with the indicated CM; G (Grow CM) and R (Ras CM). (I) Quantification of relative gap width of MDA-MB-231 cultures treated with the indicated CM. n = 3, ** p < 0.01, **** p < 0.0001.

### LCN2 upregulation is required for SASP-induced cell plasticity

Previous studies have suggested that LCN2 promotes breast cancer progression by enhancing migratory and invasive capabilities of breast cancer cells (24, 25). On account of the substantial LCN2 upregulation we detected in cells treated with senescent CM, we hypothesize that the SASP enhances aggressive breast cancer phenotypes at least in part through upregulation of LCN2. We first successfully inactivated LCN2 in MCF7 cells by CRISPR/Cas9 induced gene deletion, as evidenced by Western Blot analysis (Figure 3A, B). In agreement with the undetectable to low levels of LCN2 expressed in MCF7 cells grown in normal conditions, LCN2^-/-^ MCF7 cells did not exhibit any proliferation defects (data not shown). Strikingly, scratch assays indicated that the SASP-induced increase in migration in MCF7 was largely dependent on the presence of LCN2. Indeed, exposure to the SASP had no noticeable impact on the ability of LCN2^-/-^ MCF7 to close the gap left by the scratch even after 3 days (Figure 3C, D). We also tested the migration capabilities of LCN2^-/-^ cells via transwell assay, and consistent with the scratch assay results, exposure to the SASP did not promote migration in MCF7 LCN2^-/-^ cells (Figure 3E). To determine the transcriptional programs engaged by LCN2 in breast cancer cells exposed to the SASP, transcriptome analysis was performed on SASP-treated LCN2^+/+^ and LCN2^-/-^ MCF7 cells. Using a log2 fold change cutoff 1 and an adjusted p-value < 0.05, we confirmed LCN2 as one of the most differentially expressed genes (Figure 3F). Gene Ontology analysis indicated that pathways upregulated in LCN2^+/+^ samples include Epithelial to Mesenchymal transition and E2F targets (Figure 3G). Similarly, differentially enriched gene sets by GSEA in wild type versus LCN2^-/-^ cells, included “Epithelial to Mesenchymal Transition” as well as “MYC targets” and “TNFα Signaling via NF-κB”. Notably, these pathways have been associated with a loss of cell identity and induction of plasticity in breast cancer (26–28). Collectively, these results indicate that LCN2 is required for SASP-induced plasticity of breast cancer cells.

**Figure 3.**
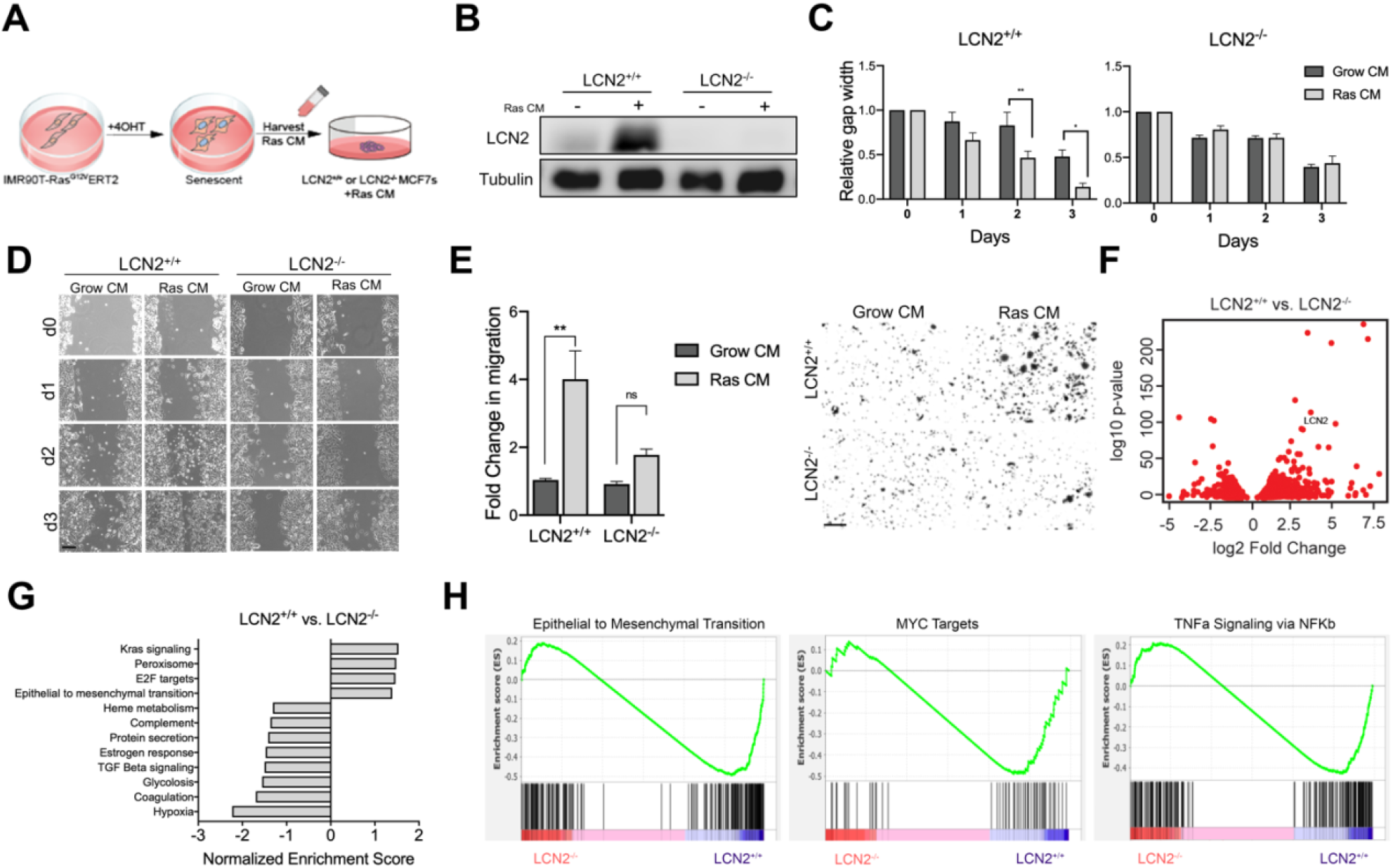
LCN2 upregulation is required for SASP-induced cell plasticity. (A) LCN2^+/+^ and LCN2^-/-^ MCF7 cells were cultured with CM from either growing or senescent cells (RasCM). (B) Western blot analysis for LCN2 expression, LCN2^+/+^ and LCN2^-/-^ MCF7 cells cultured with the indicated CM. (C, D) Representative images of scratched LCN2^+/+^ and LCN2^-/-^ MCF7 cells cultured in conditioned media (CM) from growing (Grow CM) or Ras-senescent (Ras CM) IMR90T cells for 3 days. Scale bar = 200 um. (E) Representative images and quantification of transwell migration assays of LCN2^+/+^ and LCN2^-/-^ MCF7 cells treated with the indicated CM for 2 days. Scale bar = 200 um. (F) Volcano plot depicting differentially expressed genes in LCN2^+/+^ compared to LCN2^-/-^ when treated with Ras CM. (G) Gene ontology analysis of genes that are differentially expressed in LCN2^+/+^ compared to LCN2^-/-^ when treated with Ras CM. (H) GSEA plots of pathways affected by LCN2 knockout when treated with Ras CM. n = 3, * p < 0.05, ** p < 0.01.

### SASP-induced LCN2 expression promotes breast cancer progression *in vivo*

We next sought to assess the impact of LCN2 expression on tumor progression *in vivo*. We injected Luciferase-expressing MDA-MB-231 cells into mammary fat pads of nude mice and followed tumor progression. Importantly, LCN2 status had no impact on MDA-MB-231 tumor growth (Figure 4A, B). This observation is consistent with the undetectable to low LCN2 expression levels in these cells in the absence of a pro-senescence stimulus (Figure 2H). Importantly, Western Blot analysis revealed that SASP-dependent upregulation of LCN2 levels was only transient, as removal of conditioned media from senescent fibroblasts resulted in a strong downregulation of LCN2 levels as early as 2 days (Figure 4C). Therefore, to ensure continuous SASP exposure and upregulation of LCN2 in tumors, we opted for a co injection model with senescent human fibroblasts. MDA-MB-231 cells exhibited increased LCN2-dependent proliferation and tumor progression when co-injected with senescent fibroblasts (Figure 5D-F). IHC revealed that LCN2^+/+^ tumor cells expressed reduced levels of E-cadherin and increased Ki67, compared to their LCN2^-/-^ counterparts. Apoptosis could not account for the difference in tumor size between experimental groups since LCN2^-/-^ tumors were negative for cleaved-caspase 3 (Figure 5G). These results indicate that SASP-dependent LCN2 upregulation promotes tumor cell plasticity *in vivo* and results in the development of more aggressive tumors that are highly proliferative.

**Figure 4.**
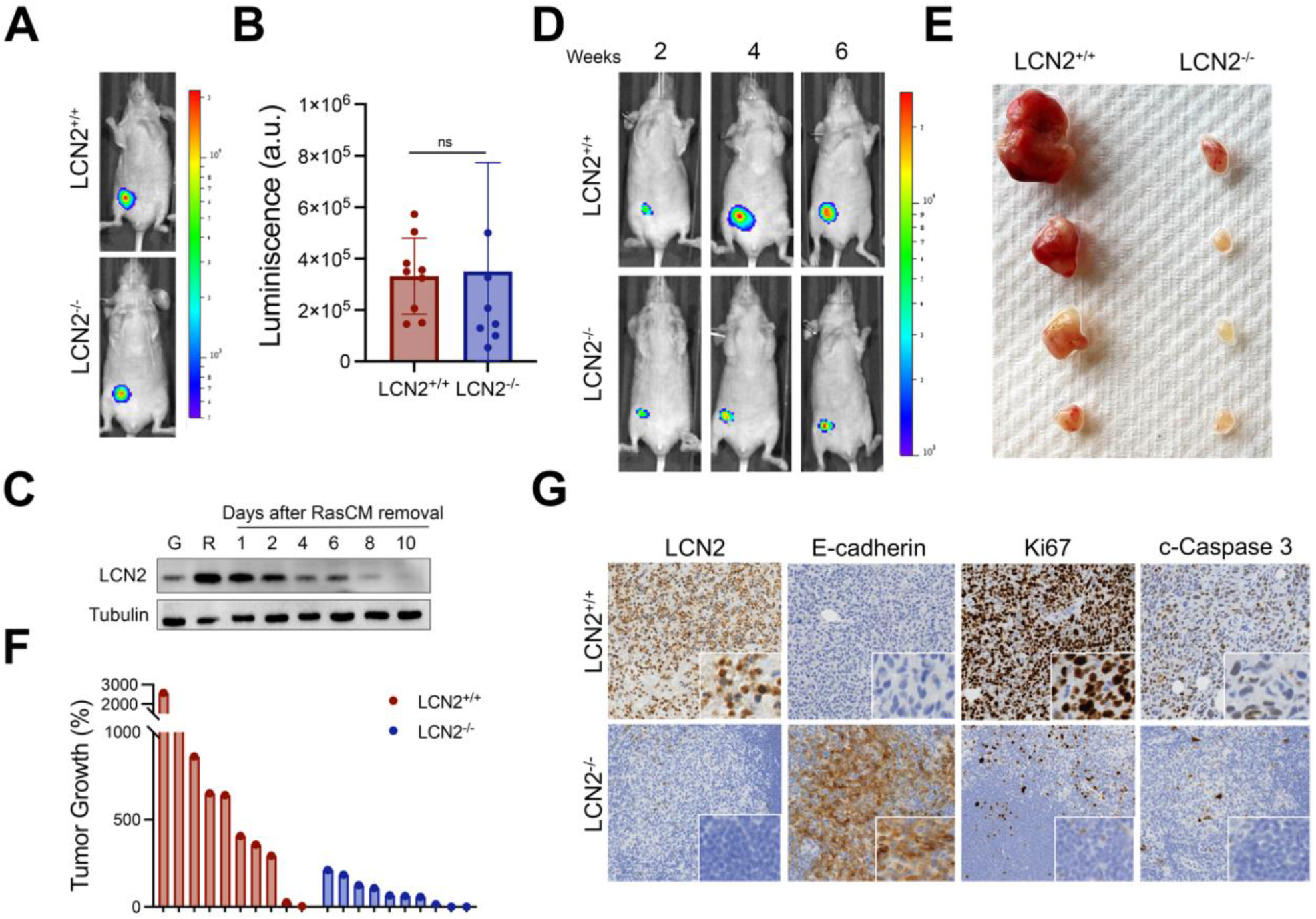
SASP-induced LCN2 expression promotes breast cancer progression *in vivo*. (A) Representative images of tumor growth progression of LCN2^+/+^ and LCN2^-/-^ MDA-MB-231 GFP-Luciferase injected into mammary fat pad of nude female mice. (B) Luminescence quantification for (A) using a PerkinElmer IVIS Spectrum system. n = 10. (C) Western blot analysis for LCN2 expression of MDA-MB-231 cells after removal or RasCM, G (Grow CM) and R (Ras CM). (D) Representative images of tumor growth progression of LCN2^+/+^ and LCN2^-/-^ MDA-MB-231 GFP-Luciferase co injected into mammary fat pad of nude female mice with 1×10^6^ senescent fibroblasts. (E) Images of dissected LCN2^+/+^ and LCN2^-/-^ tumor tissue. (F). Waterfall plot of the percentage change in tumor growth for LCN2^+/+^ and LCN2^-/-^ MDA-MB-231 GFP-Luciferase co injected with senescent fibroblasts. n = 10. (G) Representative images of IHC for LCN2, E-cad, Ki67 and c-Caspase 3 of dissected LCN2^+/+^ and LCN2^-/-^ tumors.

**Figure 5.**
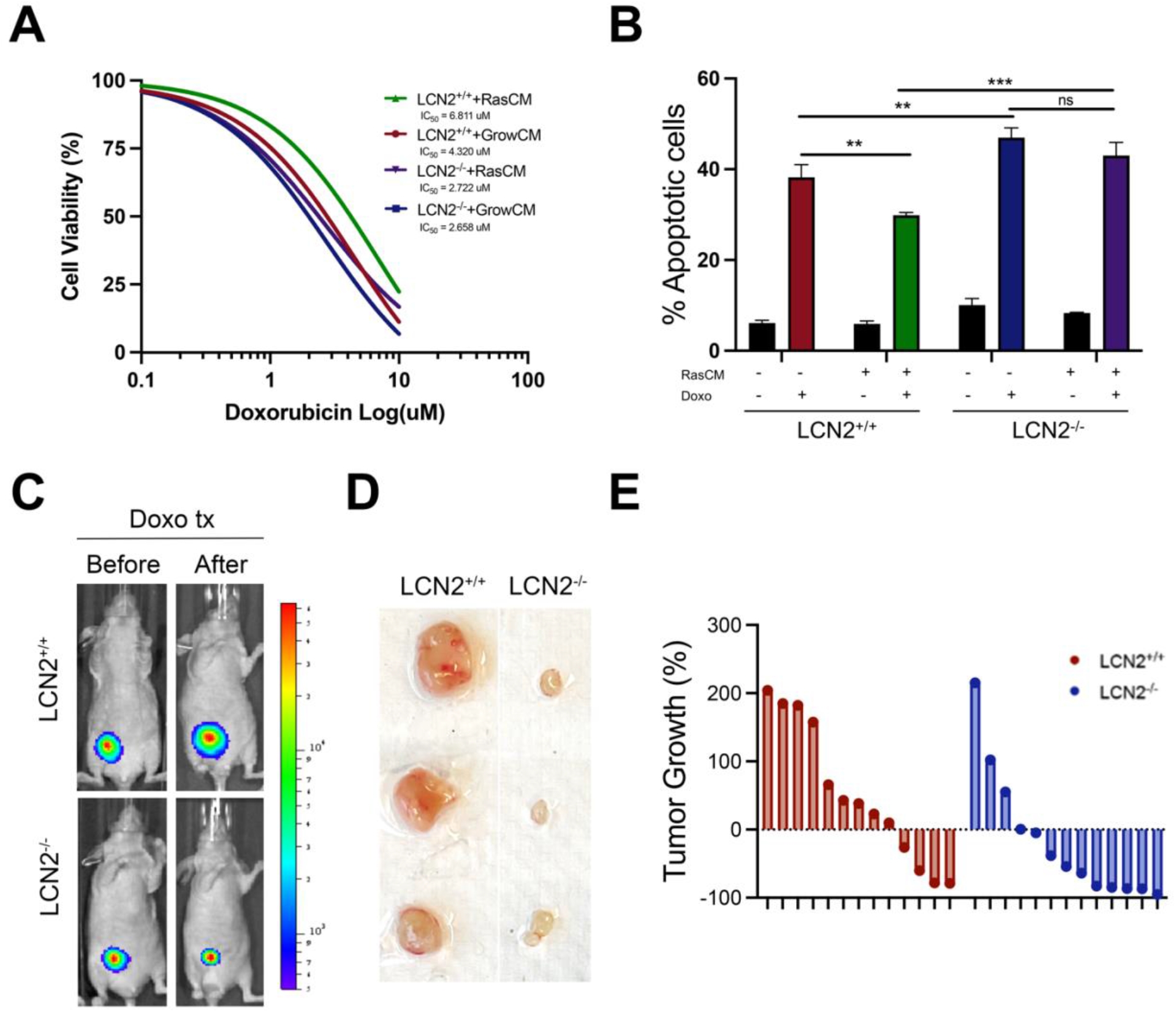
SASP-induced LCN2 protects breast cancer cells from genotoxic stress. (A) Cell viability of LCN2^+/+^ and LCN2^-/-^ MDA-MB-231 cells treated with increasing concentrations of doxorubicin and CM. (B) Annexin V quantification for LCN2^+/+^ and LCN2^-/-^ MDA-MB-231 cells treated with 4 uM doxorubicin and CM for 24 hours. n = 3, ** p < 0.01, *** p < 0.001. (C) Representative images of tumor growth progression of LCN2^+/+^ and LCN2^-/-^ MDA-MB-231 GFP-Luciferase injected into mammary fat pad of nude female mice; before and 1 week after treatment with 10 mg/kg doxorubicin. (D) Images of dissected LCN2^+/+^ and LCN2^-/-^ tumor tissue 1 week after treatment with 10 mg/kg doxorubicin. (E) Waterfall plot of the percentage change in tumor growth for LCN2^+/+^ and LCN2^-/-^ MDA-MB-231 GFP-Luciferase tumors treated with doxorubicin. n = 13.

### SASP-induced LCN2 protects breast cancer cells from genotoxic stress

Tumor cell plasticity and EMT are closely associated with therapy resistance in breast cancer (29–31). Based on the phenotypes elicited by LCN2^-/-^ breast cancer cells *in vivo*, we hypothesized that SASP-mediated LCN2 upregulation could confer breast cancer cells a chemo-protective phenotype. Indeed, cell viability and Annexin V assays indicated that LCN2^+/+^ MDA-MB-231 cells exposed to SASP were more resistant to doxorubicin than cells exposed to normal medium or LCN2^-/-^ cells (Figure 5A, B). To validate these findings *in vivo*, we injected MDA-MB-231 cells into mammary fat pads of nude mice. Once tumors were established, mice were treated with doxorubicin. While LCN2^+/+^ and LCN2^-/-^ tumors grew at a comparable rate prior to doxorubicin treatment, doxorubicin injection resulted in a dramatic sensitization of LCN2^-/-^ tumors compared to their LCN2^+/+^ counterparts (Figure 5C-E). One week after doxorubicin treatment, LCN2^+/+^ tumors were significantly larger than their LCN2^-/-^ counterparts, indicating that resistance to doxorubicin is enhanced by SASP-induced LCN2. Together, these results suggest that through the engagement of the SASP, LCN2 expression promotes detrimental cellular plasticity in breast tumors.

### LCN2 expression is induced following chemotherapy and is a poor prognostic factor in breast cancer patients

To assess the clinical relevance of the previous observations, we analyzed LCN2 levels in biopsy samples from individual breast cancer patients collected prior to or following neo-adjuvant chemotherapy treatment. Upon chemotherapy treatment, we detected an increased positivity for the SASP marker IL-6 (Figure 6A, B). Along IL-6 upregulation, samples collected after neoadjuvant chemotherapy treatment exhibited increased LCN2 positivity, while all biopsy samples collected before chemotherapy treatment displayed undetectable or low LCN2 expression (Figure 6A, C). An opposite pattern of expression was observed for the epithelial marker, E-cadherin (Figure 6A, D). We further analyzed the correlations between LCN2 levels and prognosis in breast cancer patients using publicly available expression databases. The analysis revealed that LCN2 was upregulated at the mRNA level in 125 of 2,507 patient samples (7%). Patients with increased levels of LCN2 have an inferior overall (Data not shown) and relapse-free survival compared to those with unaltered levels of LCN2 (Figure 6E). Additionally, 52.8% of patients with high LCN2 levels had received chemotherapy treatment prior to analysis, while only 18.5% of the patients with low levels of LCN2 had (Figure 6F), further indicating a correlation between chemotherapy treatment and increased LCN2 levels. These data suggest that LCN2 could be a potential prognostic biomarker for breast cancer survival.

**Figure 6.**
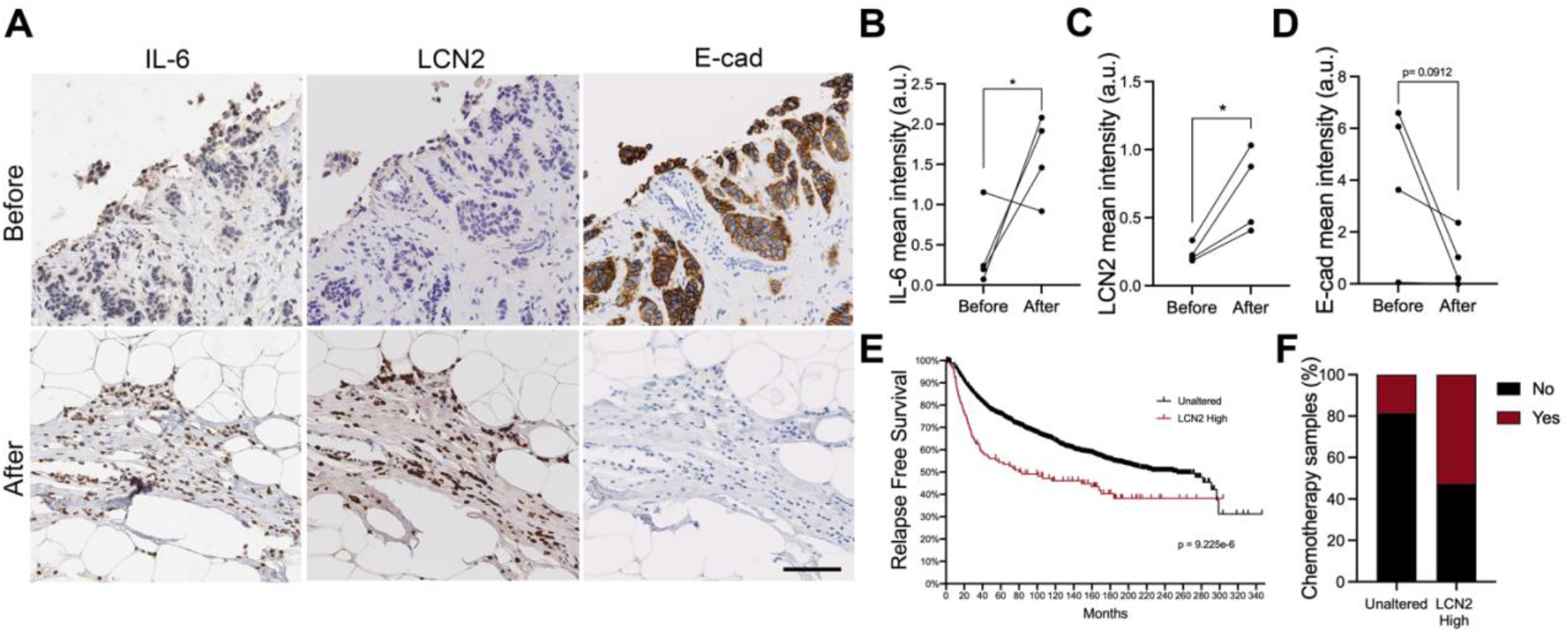
LCN2 expression is induced following chemotherapy and is a poor prognostic factor in breast cancer patients. (A) Representative images of IHC for IL-6, LCN2 and E-cad before and after neoadjuvant therapy. Scale bar = 100 um. (B) Quantification of mean intensity of IHC for IL-6 normalized by cell number per slide. (C) Quantification of mean intensity of IHC for LCN2 normalized by cell number per slide. (D) Quantification of mean intensity of IHC for E-cad normalized by cell number per slide. (E) Kaplan-Meier plots showing progression-free survival of patients with unaltered LCN2 (median, 248.95 months) and patients with high LCN2 levels (median, 83.32 months). (F) Plot showing the percentage of breast cancer patients that received chemotherapy and their LCN2 status. * p < 0.05.

## Discussion

Cellular plasticity promotes tumor progression in breast cancer, through metastatic spread and resistance to conventional therapies. The molecular pathways that contribute to the plasticity of breast cancer cells, including EMT and dedifferentiation, are currently being uncovered. However, the stimuli leading to the acquisition of a cellular plastic phenotype within an established tumor remain largely unknown. We report here that exposure to the SASP induces breast cancer cell plasticity in an LCN2-dependent manner.

Our results are consistent with previous findings showing that breast cancer cells exposed to the SASP exhibit epithelial cell scattering and reduced cell-cell adhesions, features of EMT, and increased expression of stemness markers (32, 33). Strikingly, we demonstrate that LCN2 upregulation is critical for the engagement of an EMT-associated expression program in breast cancer cells upon SASP exposure. Furthermore, our results indicate that different molecular subtypes of breast cancer cells are equally susceptible to SASP-induced LCN2 upregulation, and the subsequent increase in cell migration. Such observation suggests that a large proportion of breast tumors could acquire more aggressive traits when surrounded by senescent cells. In the context of breast cancer, the source of the SASP can be several-fold: replicatively exhausted cells that accumulate in the organism with age secrete SASP factors that can reach tumor cells. Another source of the SASP may stem from pre-cancerous or cancerous cells themselves with constitutively active drivers of mitogenic signals, resulting in hyper-proliferation and fork collapse, known as oncogene-induced senescence (OIS). Finally, exposure to genotoxic therapies, including radiation therapy or chemotherapy, can drive both normal and tumor cells into senescence, in a process termed Therapy-Induced Senescence (TIS), leading to general and local SASP production. TIS has emerged as a novel functional target to improve cancer therapy (34). However, accumulating evidence indicates that senescent cancer cells are capable of reentering the cell cycle and promoting tumor relapse and metastasis. Recent findings show that a senescence-like population of chemotherapy-resilient cells is capable of initiating cancer recurrence by increasing their stemness potential (35–37).

Previous studies have linked LCN2 and EMT or metastasis of breast cancer cells (38, 39), but the mechanisms involved remain unclear. Downregulation of the estrogen receptor ERα induces expression of the transcription factor Slug, driving LCN2-induced EMT. Our transcriptome analyses indicate that ERα is indeed downregulated and Slug is upregulated in MCF7 cells treated with senescent CM compared to those treated with growing CM (data not shown). Studies have demonstrated that ERα signaling helps maintain the epithelial phenotype through inhibition of Snail activity (40). However, based on our demonstration that SASP-induced LCN2 promotes migration in ER-breast cancer cells, it is unlikely that LCN2 mediates its effects on breast cancer plasticity through the ERα pathway exclusively. Alternatively, LCN2 could drive EMT through its interaction with MMP9, reducing E-cadherin expression levels on the cell surface (41). Further experiments will be necessary to elucidate the mechanism employed by LCN2 to promote EMT in breast cancer cells.

Lipocalin-2 (LCN2) was first characterized as an iron-binding protein, sequestering it from Gram-negative bacteria and inhibiting their proliferation. LCN2 iron-binding sequestration properties have also been implicated in the etiology of Leptomeningeal Metastasis (LM) (42). Indeed, LCN2 upregulation allows metastatic cancer cells to thrive in an iron-limiting environment such as the cerebrospinal fluid. Accordingly, the administration of iron chelators slows cancer progression in a mouse model of LM (42). Iron facilitates tumorigenesis by driving cell proliferation (43). Therefore, LCN2 upregulation could result in an increased ability to sequester iron leading to an aggressive tumor growth in breast cancer cells (44).

The experiments presented and conclusions drawn here focus on the cancer cell autonomous impact of SASP-induced LCN2 upregulation. However, based on the pleiotropic impact of iron metabolism, it is also likely that LCN2 modulates the anti-tumor immune response *in vivo*. Indeed, LCN2 has the ability to upregulate human leukocyte antigen G (HLA-G), which can promote tumor immune escape in mouse models (45). Furthermore, by conferring cancer cells with the ability to outcompete macrophages for iron, LCN2 may also contribute to the generation of a tumor microenvironment that promotes tumor growth, a possibility that remains to be investigated. For these reasons, therapeutic inhibition of LCN2 could provide a therapeutic relief for patients with breast tumors that are poised to undergo plasticity, for example following exposure to genotoxic therapies.

## Materials and Methods

### Cells

IMR90 primary lung embryonic fibroblasts expressing hTERT (IMR90Ts) were obtained from S. Smith (NYU School of Medicine, New York, NY). Cells were cultured in MEM (Corning) supplemented with 10% fetal bovine serum and 1% penicillin-streptomycin (Cellgro). MCF7, SKBR3, Hs578t and T47D cells were obtained from R. Possemato (NYU School of Medicine, New York, NY). MDA-MB-231 cells were obtained from E. Hernando (NYU School of Medicine, New York, NY). BT474 cells were obtained from B. Neel (NYU School of Medicine, New York, NY). Breast cancer cells (except T47D) were cultured in DMEM (Corning) supplemented with 10% FBS and 1% penicillin/streptomycin. T47D cells were cultured in RPM1 1640 media (Corning) supplemented with 10% FBS and 1% penicillin/streptomycin. HEK293T cells (ATCC) were used to generate retro- and lentiviruses and were cultured in DMEM (Corning), supplemented with 10% donor calf serum and 1% penicillin-streptomycin. IMR90T cells were maintained in 6% O_2_ and 5% CO_2_ at 37°C, while 293T and breast cancer cells were maintained in 5% CO_2_ at 37°C.

### Senescence induction and condition media (CM) harvest

For Ras-induced senescence, IMR90T-Ras-ERT2 were treated with 200 nM tamoxifen (Sigma) continuously for 10 days. Fresh media and tamoxifen were added every 2-3 days. Cells were treated with an equal volume of ethanol as control. On day 8 of Ras induction, media was replaced with serum-free media and harvested after 48 h. For etoposide-induced senescence, cells were treated with 50 uM etoposide (Sigma) or an equal volume of DMSO as a control for 48 hours. Etoposide-containing medium was then replaced by normal culture media for 5 days. 7 days after etoposide treatment, media was replaced with serum-free media and harvested after 48 h. Conditioned media was aliquoted and flash frozen in liquid nitrogen before storing at −80 C.

### Scratch assays

MCF7 or MDA-MB-231 cells were grown in 6-well plates until confluent. A P200 tip was used to create vertical scratches. Media was then changed to media containing 1% FBS supplemented by CM from growing or senescent cells. Amount of CM added was normalized to cell number. CM was replaced every day. Pictures of scratches per well were taken each day for 3 days. Using Adobe Photoshop, the gap width was measured as the number of pixels that comprise the gap. The data is presented as relative gap width compared to gap width of each sample at day 0.

### Transwell migration assays

MCF7 cells were exposed to 1-3 mL CM from growing or senescent cells. Fresh CM was added every day for a total of 2 days. Cells were then trypsinized and counted. A total of 50,000 cells in 100 uL serum-free DMEM were seeded on top of transwells containing 8 um pores for use in 24-well plates. 1 mL of DMEM supplemented with 20% FBS was added to the bottom of the wells and cells were allowed to migrate for 48 h. Transwells were then washed in PBS and fixed in 1% glutaraldehyde in PBS for 20 minutes, and crystal violet for 30 min rocking. Values are expressed as fold change in the number of cells compared to the number of cells cultured in the corresponding growing CM.

### Immunofluorescence

MCF7 cells were treated with CM from growing or senescent cells. CM was added every day for 2 days. Cells were then plated on coverslips before being fixed in 4% paraformaldehyde (SCBT) for 10 min at RT. Cells were permeabilized with cold 0.1% Triton X-100 in PBS for 5 min and blocked with 20% donkey serum in PBST for 15 min. Cells were incubated with mouse anti-E-cadherin (Millipore, MAB1199) at 1:200 dilution in blocking solution at 37°C for 1 h. Cells were then washed and incubated in Cy3-conjugated donkey anti-mouse IgG (Jackson Immunoresearch) for 1 hour at RT. Cells were mounted with mounting medium containing Dapi (Vectashield). Slides were examined on a Zeiss AxioImager A2 microscope. A total of 100 cells per cover-slip were counted. Amount E-cadherin staining was quantified as number of red pixels per Dapi-positive cells using Image J to calculate pixels.

### Transcriptomics analysis

MCF7 cells were cultured in CM from growing or Ras-induced senescent cells for 2 days. MCF7 LCN2^+/+^ and LCN2^-/-^ cells were cultured in CM from Ras-induced senescent cells for 2 days. RNA quality assessment, library preparation and sequencing were performed by the NYU School of Medicine Genome Technology Center or by Genewiz. Strand-specific libraries were prepared using the TruSeq RNA library Prep kit, and libraries were sequenced on an Illumina HiSeq2500 using 50-bp paired-end reads. Sequences were mapped to the hg10 genome, and analysis was done as previously described (Proudhon, 2016).

### Real-Time PCR

Total RNA was extracted using TRIzol (Life Technologies) according to manufacturer’s instructions. 1 ug of total RNA was used to generate cDNA with oligo dT. Real-time qPCR was performed using Maxima SYBR Green (Fisher Scientific) and samples were run on a Bio-Rad Cycler MyiQ. The following sets of primer were used: *hTubulin* forward 5’-cttcgtctccgccatcag-3’, reverse 5’-ttgccaatctggacacca-3’, *hLCN2* forward 5’-gaagtctgactactggatcagga-3’, reverse 5’-accactcggacgaggtaact-3’, *mGAPDH* forward 5’-cacggcaaattcaacggcacagtc-3’, reverse 5’-acccgtttggctccacccttca-3’, *mLCN2* forward 5’-gacttccggagcgatcagtt-3’, reverse 5’-ttctgatccagtagcgacagc-3’.

### CRISPR/Cas 9 editing

sgRNAs were cloned into lentiCRISPR v2 (Addgene) and 48 hours after infection, cells were selected with puromycin (1 ug/mL) for at least 4 days before plating single cell clones. The following sgRNA’s were used for hLCN2: forward 5’-caccgaagtggtatgtggtag-3’, reverse 5’-aaacggcctaccacataccacttc-3’ and forward 5’-caccgtggtggcatacatctt-3’, reverse 5’-aaacgcaaaagatgtatgccacca-3’.

### Protein extraction and Western blotting

Cells were lysed in 1× RIPA buffer (1% NP-40, 0.1% SDS, 50 mM Tris-HCl, pH 7.4, 150 mM NaCl, 0.5% sodium deoxycholate, 1 mM EDTA), 0.5 μM DTT, 25 mM NaF, 1 mM sodium vanadate, 1 mM PMSF, and cOmplete protease cocktail inhibitor (Sigma). Samples were resolved by SDS-PAGE and analyzed by standard western blotting techniques. The following primary antibodies were used: mouse anti-tubulin (Sigma T9026) at 1:2000 dilution, goat anti-LCN2 (R&D AF1747) at 0.2 ug/mL, mouse anti-vinculin (Sigma V9131) at 1:1000 dilution and rabbit anti-EGFR (Cell Signaling 4267) at 1:1000 dilution.

### Bioluminescence

For in vivo luminescence of Luciferase, mice were injected i.p. with of 150 mg of D-Luciferin (ThermoFisher) per kg of body weight. Fifteen minutes later, the mice were anesthetized with isoflurane and luminescence was measured with a PerkinElmer IVIS Spectrum system.

### MDA-MB-231 co-injection with senescent fibroblasts

2.5×10^5^ LCN2^+/+^ or LCN2^-/-^ MDA-MB-231 GFP-Luciferase cells were pretreated with RasCM for 48 hours before being injected into the inguinal mammary fat pad of 4-6 weeks old female nude mice with 1×10^6^ senescent IMR90T cells in 50% Matrigel (Corning).

### Immunohistochemical staining

Immunohistochemical staining was performed at the NYU Langone Experimental Pathology Research Laboratory as previously described (Rielland, 2014). The following antibodies were used: goat anti-LCN2 (R&D AF1747), E-cadherin (Cell Signal, 3195T) IL-6 (Santa Cruz Biotechnology, sc-1265).

### Cell viability assay

400, 000 cells were plated in triplicate in 12-well plates and allowed to adhere overnight. Cells were then treated with increasing concentrations of doxorubicin and CM for 24 hours. Wells were washed with PBS and cells were fixed with 2% glutaraldehyde in PBS for 15 minutes. Cells were then stained with crystal violet (0.1% in 10% ethanol) for 30 minutes. After washing and drying, cells were distained in 10% acetic acid for 15 minutes. Optical density (OD) was measured at 595 nm absorbance.

### Annexin V staining

Treated and untreated cells were collected, without washing to collect all floating cells, centrifuged at 1300 rpm for 3 minutes. After discarding supernatant, 5 uL of Annexin V (BioLegend) was added to cells resuspended in 200 uL of binding buffer (BioLegend). Cells were then incubated for 30 minutes at room temperature in the dark before centrifuging them again. Cells were resuspended in 200 uL of binding buffer after discarding the supernatant and analyzed via flow cytometry using a FACSCalibur Flow Cytometer (BD).

### MDA-MB-231 tumors and doxorubicin treatment

1×10^5^ LCN2^+/+^ or LCN2^-/-^ MDA-MB-231 GFP-Luciferase cells in 50% Matrigel (Corning) were injected into the inguinal mammary fat pad of 4-6 weeks old female nude mice. Once tumors were established, mice were treated with a single dose of 10 mg/kg of doxorubicin (Sigma).

### Statistical Analysis

Results were analyzed using GraphPad Prism software. Values were subjected to unpaired two-tailed t-tests, multiple t-tests, one-way ANOVA followed by Dunnett’s multiple comparisons test, or two-way ANOVA followed by Tukey’s multiple comparisons test. Data is presented as means ± SEM.

### Study Approval

Animal work and human subject work were performed accordingly to approved protocols from the NYU School of Medicine’s IACUC and IRB. Written informed consent was obtained from each human participant before study procedure.

## Acknowledgements

The authors sincerely thank all members of the David lab for helpful discussions during the preparation of this manuscript. We wish to acknowledge the NYU Genome Technology Center for help with RNA sequencing (RNA-seq). We thank Dr. Richard Possemato (NYU School of Medicine), Dr. Eva Hernando (NYU School of Medicine), Dr. Benjamin Neel (NYU School of Medicine) and Dr. Judith Campisi (Buck Institute for Research on Aging) as well as members of their labs for the generous gift or reagents and plasmids and for helpful discussions. This work was funded by NIH/NCI (CA246416) [GD], NYS DoH (C36617GG) [GD] and the NYSTEM Institutional Training Grant (C322560GG) [JMV].

## Author contributions

The authors confirm contribution to the paper as follows: LL initiated the study under supervision of GD and contributed to figures 1 and 2 of the manuscript. JMV designed, performed and analyzed data from the experiments presented in this manuscript. TMN contributed to sample preparation. UD provided patient samples and analyzed the data. GD designed the study and supervised the research. JMV and GD wrote the manuscript. All authors reviewed the results and approved final version of the manuscript.

